# Hypertonic saline solution inhibits SARS-CoV-2 in vitro assay

**DOI:** 10.1101/2020.08.04.235549

**Authors:** Rafael R. G. Machado, Talita Glaser, Danielle B. Araujo, Lyvia Lintzmaier Petiz, Danielle B. L. Oliveira, Giuliana S. Durigon, Alessandra L. Leal, João Renato R. Pinho, Luis Carlos S. Ferreira, Henning Ulrich, Edison L. Durigon, Cristiane R. Guzzo

## Abstract

We are facing an unprecedented global health crisis caused by severe acute respiratory syndrome coronavirus 2 (SARS-CoV-2). At this date more than 680 thousand people have died due to coronavirus disease 2019 (COVID-19). Unfortunately, until now no effective treatment to combat the virus and vaccine are available. We performed experiments to test if hypertonic saline solution is able to inhibit virus replication *in vitro*. Our data shows that 260 mM NaCl (1.5%) inhibits 100% SARS-CoV-2 replication in Vero cells. Furthermore, our results suggest that the virus replication inhibition is due to an intracellular mechanism and not due to the dissociation between spike SARS-CoV-2 protein and its human receptor angiotensin-converting enzyme 2 interaction. NaCl depolarizes the plasma membrane supposedly associated with the inhibition of the SARS-CoV-2 life cycle. This observation could lead to simple, safe and low cost interventions at various stages of COVID-19 treatment, improving the prognosis of infected patients, thereby mitigating the social and economic costs of the pandemic.

## Introduction

The world is facing a pandemic situation due to the pathogenic SARS-COV-2 (Severe Acute Respiratory Syndrome Coronavirus 2)^1,2^, a zoonotic virus, that spread from China to many countries around the world in less than 3 months. So far there were more than 17 million confirmed patients in 213 countries, and more than 680,000 deaths due to COVID-19 disease, according to WHO data as of July 26, 2020 (https://covid19.who.int). Few medical procedures are available, which could reduce viral load, decreasing patient hospitalization time, virus dissemination and the number of patients with severe symptoms.

SARS-CoV-2 uses a surface glycosylated spike (S) protein to bind human angiotensin-converting enzyme 2 (ACE-2)^3^, CD147^4^ and sialic acid receptors^5^ and to develop coronavirus disease (COVID-19). In the case of SARS-CoV, that shares 76% of identity with SARS-CoV-2 S protein, it has been suggested that the virus may also bind to host cells through DC-SIGN or L-SIGN receptors^6,7^. SARS-CoV-2 and ACE-2 interaction is essential to mediate virus infection^8^. The S protein has two main subunits, the S1 subunit involved in binding to the host cellular receptor, and the S2 subunit that mediates fusion of the viral and cellular membranes. Initially, the homo-trimeric S protein via its receptor binding domain (RBD) located at the S1 subunit, binds to human ACE-2 (hACE-2), causing a conformational change in the S1 and S2 subunits leading to the fusion of virus membrane with the host cell membrane^9,10^. The SARS-CoV-2 S protein structure has been recently solved in the apo form^11^ and in complex with ACE-2 protein^12^. Interestingly, the interface between the RBD of S protein with hACE-2 is highly polar, involving charged residues located mainly at the hACE-2 interface and polar residues at the RBD interface^12^. These ionic interactions could in principle be affected by increasing the ionic strength and could be used to inhibit the interaction of the virus with the host cell. Moreover, the potential antiviral activity, *in vitro*, of sodium chloride (NaCl) has been already studied for RNA viruses, such as mengovirus^13^, respiratory syncytial virus, influenza A virus, human coronavirus 229E and coxsackievirus B3, and also for DNA virus, like herpes simplex virus-1 and murine gammaherpesvirus 68^14^. Furthermore, a randomised controlled clinical trial studying the effectiveness of the treatment for viral upper respiratory tract infection, using hypertonic saline nasal irrigation and gargle (HSNIG)^15^, observed a decrease in the duration of illness, over-the-counter medications use, transmission within household contacts and viral shedding^15^. However, the molecular details of the antiviral NaCl activity remains unresolved.

In order to test our initial hypothesis, we performed different assays to determine if different concentrations of NaCl affect the SARS-CoV-2 replication, an RNA enveloped virus, when cultured in monkey kidney epithelial cells (Vero CCL-81), and we also evaluated the effect of NaCl in the membrane potential.

## Results

### Effect of NaCl in vitro SARS-CoV-2 replication

We performed experiments to determine if hypertonic saline solution is able to inhibit SARS-CoV-2 replication *in vitro*. We measured the effects of increasing concentrations of NaCl (135, 160, 185, 210, 235, 260 and 285 mM equivalent to 0.8, 0.9, 1.1, 1.2, 1.4, 1.5, 1.7%, respectively) on cell viability and their antiviral activity on SARS-CoV-2. Efficacies were evaluated by quantification of viral copy numbers in the cell supernatant via quantitative real-time RT-PCR (RT-qPCR) and confirmed with visualization of cytopathic effect by optical microscopy at 72□hours post infection (h.p.i.). Our data shows that 210 mM NaCl (1.2%) was sufficient to inhibit the virus replication by 90% (**Fig. 1a**), achieving 100% of inhibition on 260mM (1.5%). Next, to determine which stage of virus replication was affected by the NaCl, we evaluated if viral inhibition was a direct effect of NaCl on the virus particles, for that SARS-CoV-2 was pre-incubated with media and increasing concentrations of NaCl for 1 hour before absorption. Virus pre-exposure to NaCl did not affect viral replication at any concentration of NaCl tested (**Fig. 1a**, VPI curve). The same pattern of absence of inhibition was observed when we treated the cells one hour before infection, analyzing the adsorption process, which consists of the interaction of virus with cell receptors (**Fig. 1a**, AD curve). On the other hand, significant inhibition of viral replication (up to 50%) was seen when as little as 160 mM of NaCl was available during virus replication alone (**Fig. 1a**, p = 0.032, PI curve) or full-time, during adsorption and replication (**Fig. 1a**, p = 0.030, FT curve). There was no statistically significant difference between post-infection (PI) and adsorption plus post-infection (FT) treatments (p = 0.985). These data together suggest that SARS-CoV-2 inhibition in the presence of NaCl was an intracellular mechanism and was not due to the dissociation of spike SARS-CoV-2 and human angiotensin-converting enzyme 2 (ACE-2) complexes.

**Fig.1:**
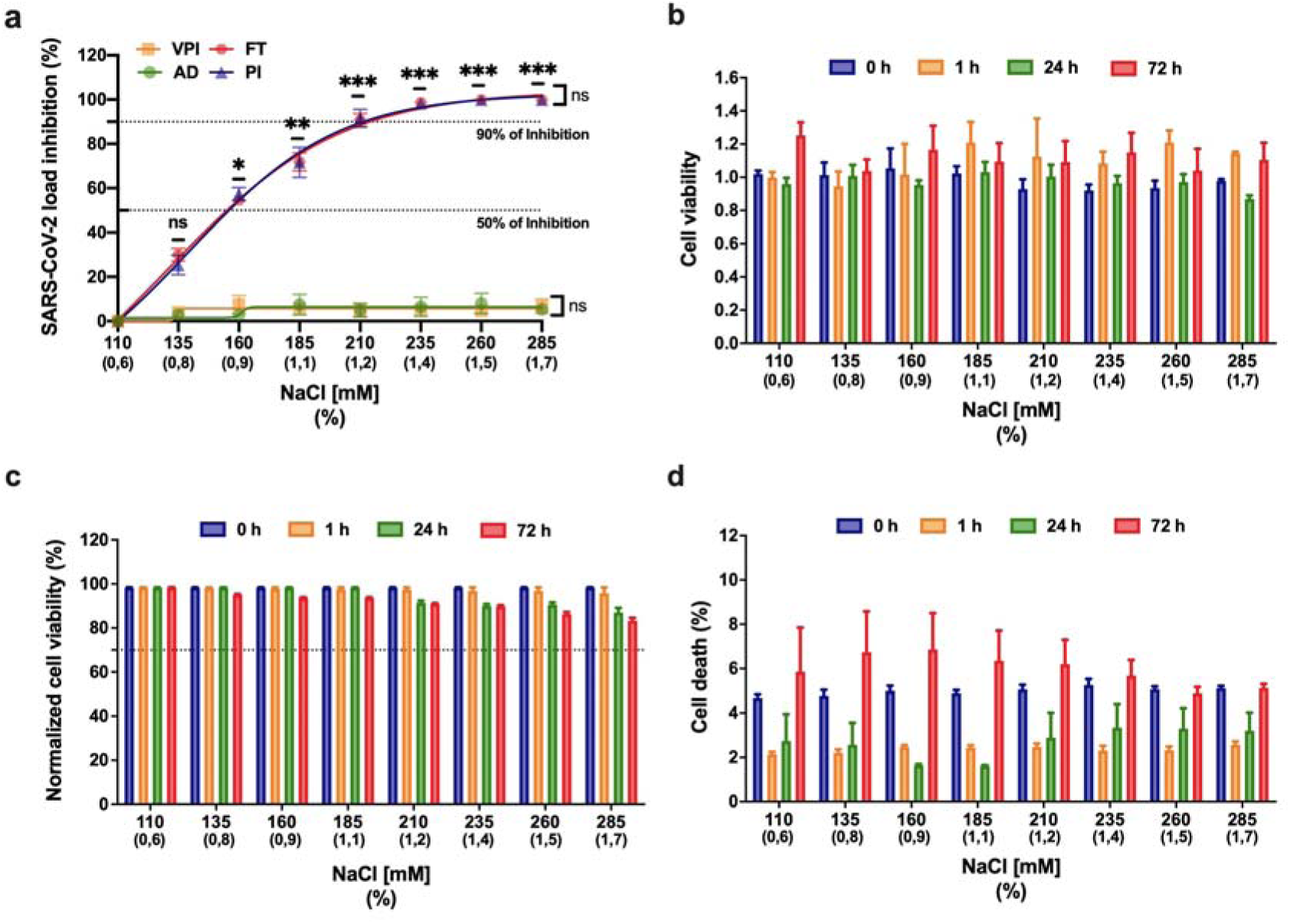
Antiviral activity of NaCl against SARS-CoV-2 *in vitro assay*. **a:** The left *Y*-axis of the graph represents percentage inhibition of virus load in cellular supernatant. Four different NaCl time-of-addition were evaluated, compromising the virus pre-incubation (VPI), absorption (AD), post-infection (PI) and adsorption plus post-infection (FT). Error bars indicate the standard error of the mean of three independent experiments carried out in triplicate each one.*p<0.05, **p<0.005 and ***p<0.0005 when compared to 110 mM NaCl. Vero CCL-81 cells were not significantly impaired in the presence of NaCl. Cells were treated with increasing concentrations of NaCl (110mM up to 285mM). Zero, 1, 24 and 72 hours post-treatment cellular viability was determined. **b:** AlamarBlue™ Cell Viability Reagent (Thermo Fisher Scientific). **c:** quantifying the LDH released in culture supernatant from cells with damaged membranes, using the CytoTox 96® Non-Radioactive Cytotoxicity Assay (Promega Corp., Madison, USA). Cell viability was normalised to untreated cells (110 mM NaCl). **d:** percentage of dead cells stained with Hoechst 33342. Viability below 70% (cell death more than 30 %) was considered evidence of cytotoxicity. Error bars represent the mean ± SEM of three independent experiments carried out in triplicate.

The antiviral activity of NaCl was not caused by cytotoxicity in the Vero cell line, as determined by the AlamarBlue™ Cell Viability Reagent (**Fig. 1b**) and LDH assay (**Fig. 1c**). Furthermore, we performed a cell death experiment to confirm the integrity of cells after being exposed to NaCl in the same conditions performed with the presence of the virus. We observed less than 8% of death at all concentrations tested (**Fig. 1d**). Related to NaCl cytotoxicity, we observed less than 20% of cytotoxicity (cell viability up to 80%) at all concentrations tested (**Fig 1b, 1c** and **1d**).

### NaCl induces cell membrane depolarization in a dose-dependent manner

Additionally, we conducted experiments to measure the hyperosmotic stress effect on cell membrane. Vero cells were incubated with different NaCl concentrations (135, 160, 185, 210, 235, 260 and 285 mM equivalent to 0.8, 0.9, 1.1, 1.2, 1.4, 1.5, 1.7%, respectively), and cell membrane potential was measured at 1, 24 and 72h after incubation (**Fig. 2a**). Results are indicated as relative fluorescence units (RFU), whose increased values refers to membrane depolarization. Cell membranes showed a rapid depolarization upon stimulation with increasing NaCl concentration (**Fig. 2b**). Depolarization was dose-dependent, with statistically difference starting at 160 mM of NaCl. The results of the 24 hours after incubation showed that the depolarization of the membrane continues to increase with time, since the values are greater than those at the time of 1 hour. Membrane potential tended to return to control values at 72h, indicating that Vero cells were partially able to recover their resting membrane potential.

**Fig.2:**
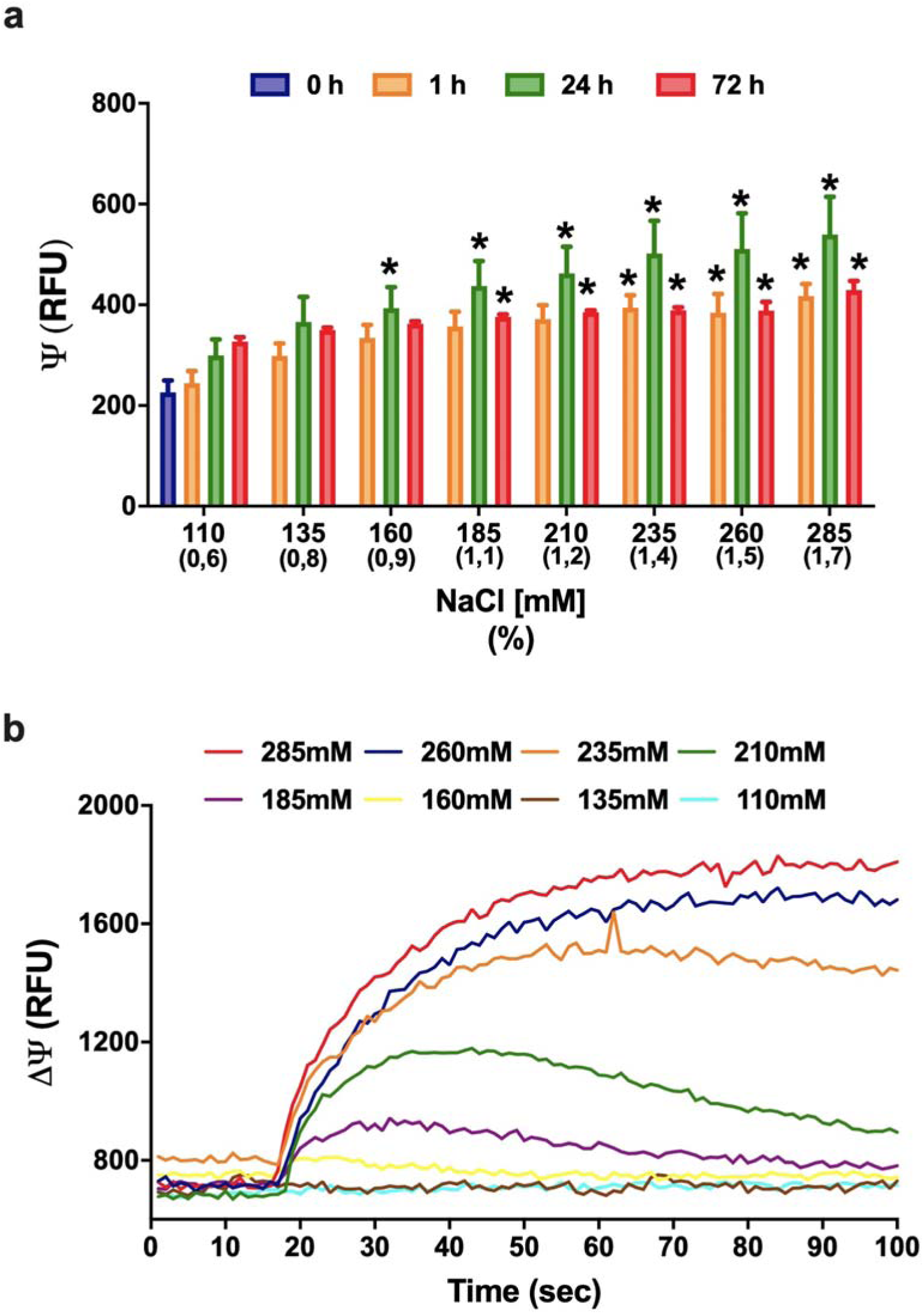
Vero cells membrane depolarization by increasing concentrations of NaCl. **a**: Membrane potential indicated as relative fluorescence units (RFU) in Vero cells after 0, 1, 24 and 72 hours post-treatment with increasing concentrations of NaCl (110 up to 285 mM) as determined by microfluorimetry. The data are representative of three independent experiments and shown as mean values ± SEM; one-way ANOVA (*p ≤ 0.05). The increasing NaCl concentration caused an immediate membrane depolarization that was maximal after 24h and tends to return to control values after 72h of treatment. **b**: The increasing NaCl concentration causes an immediate membrane depolarization in NaCl concentrations above 160 mM. The depolarization for cells treated with up to 210 mM NaCl tends to restore to the resting point after 70 seconds.

## Discussion

Our data show the efficiency of hypertonic NaCl solutions in blocking SARS-CoV-2 replication in Vero cells. This data suggests that hypertonic solution can be used as a prophylaxis and an alternative treatment for COVID-19 patients. Nevertheless, clinical trials must be done to prove the efficacy of the treatment in humans. In order to understand the reason to explain the virus inhibition by NaCl, we performed membrane potential assays that also show clearly a direct relation with membrane depolarization. The SARS-CoV-2 inhibition assays were performed using monkey kidney cells and as lung epithelial cells, may be heavily infected by SARS-CoV-2^16^ and are capable of undergoing endo- and exocytosis, being therefore a good model cell to study the SARS-CoV-2 life cycle. Vesicle formation is important for kidney and lung cells as responses to alteration of osmosis, regulated by aquaporin channels^17^, as well as for vesicle transfer between cells, which has been suggested to contribute to lung disease and possibly to the propagation of the SARS-CoV-2 virus into neighboring cells^18^.

We postulate here the importance of a hyperosmotic response by high extracellular salt concentration on the expression of aquaporin channels, which would limit endocytosis and thereby inhibit entrance of SARS-CoV-2 into cells. Endocytic, clathrin-independent, lipid-raft involving virus entry has been described for SARS-CoV ^19^, supporting our proposed mechanism of endocytosis participation in SARS-CoV-2 entry. During SARS-CoV-2 infection, pulmonary edema and cytokine storm are responsible for the most lethal scenarios^20^. The increased amount of liquid blocking the airway at the alveoli causes extreme injury, and a scar fibrotic tissue substitutes the lesioned tissue (**Fig. 3a**). Along the embryonic pulmonary development, the organ is full of liquid, which is removed mainly by the action of sodium channels such as epithelial sodium (Na^+^) channels (ENaC) that are sensitive to amiloride ^21–24^. These ENaC are expressed mainly by epithelial cells of many organs, such as kidney, lungs, colon, skin, and by some neurons in the brain^21^. In adulthood ENaC usually controls the concentration of Na^+^ ions in the extracellular environment. Therefore, the channels are extremely important for the organism water balance, by controlling the reabsorption of Na^+^ in kidneys, sweat and colon. In this way, the activity of ENaC can influence blood pressure and the renin-angiotensin system, as well as the thickness and dryness of the airway^21^. Interestingly, this channel is also involved in the taste sensation and its improper functioning could be related to loss of taste ^21^.

**Fig 3:**
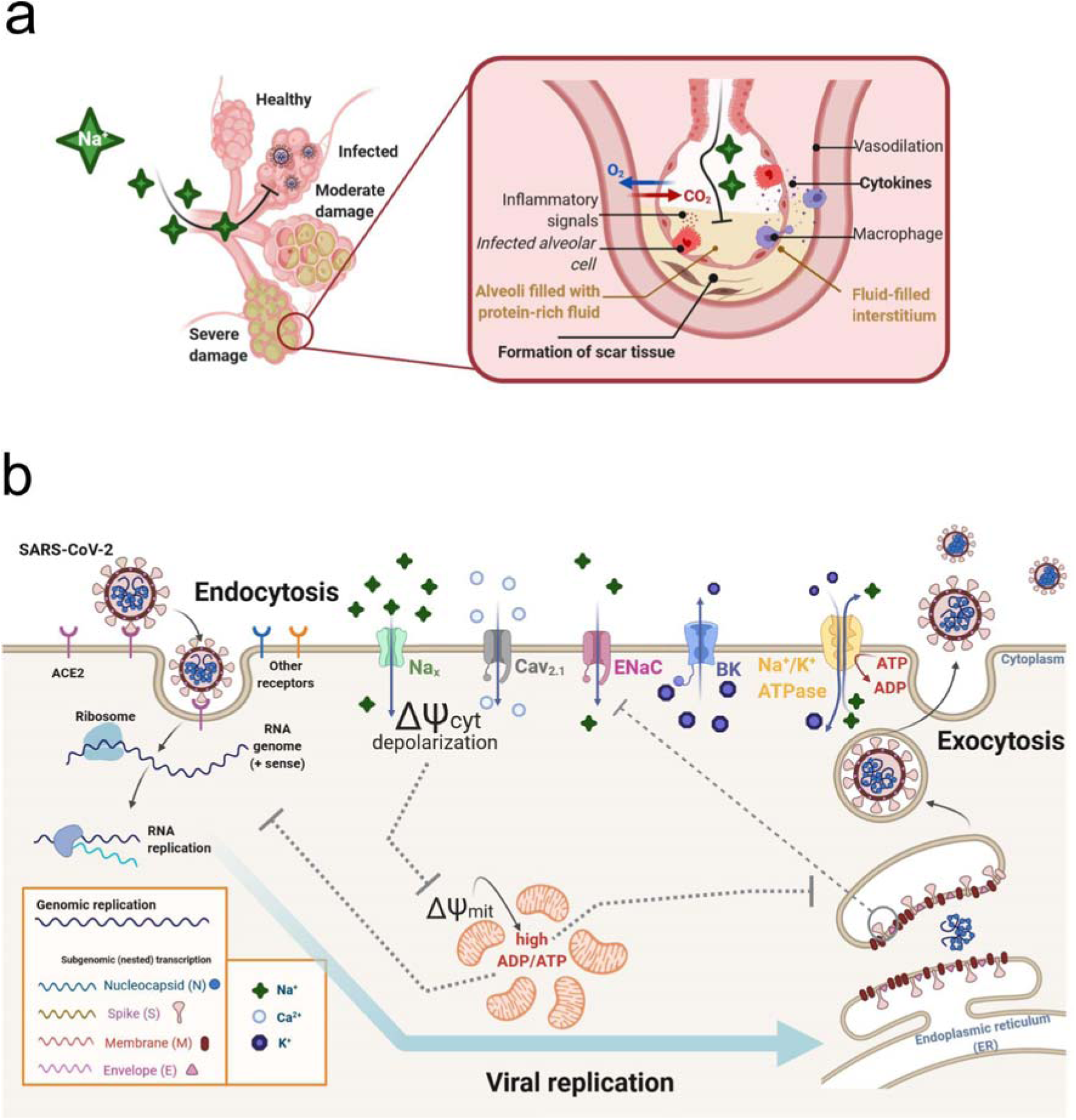
Possible mechanisms involving NaCl hyperosmotic stress and SARS-CoV-2 inhibition in Vero cells. **a:** Illustration of SARS-CoV-2 alveoli infection, which can lead to pulmonary edema, high levels of cytokines and tissue damage. **b:** Hypothesis for a possible mechanism involving the effect of NaCl in the inhibition of SARS-CoV-2 replication in Vero cells. Hypertonic saline solution causes membrane depolarization and an overflow of Na^+^ in cells, followed by increased cytosolic Ca^2+^ causing low energy state (high ADP/ATP ratio), leading to impaired virus replication.. Hyperosmotic extracellular NaCl concentrations activate Na_x_ channels, a sodium-sensitive (but not voltage-sensitive) channel, which is critically involved in body-fluid homeostasis. High cytoplasmic Na^+^ can recruit epithelial sodium (Na^+^) channels (ENaC), further increasing intracellular Na^+^ concentration, causing cell membrane to depolarize and to open voltage-gated calcium channels Cav2.1. The membrane depolarization and inward Ca^2+^ current would cause impairment in mitochondrial function, which combined to Na^+^/K^+^ ATPase high activity – due to cell attempt to restore rest membrane potential – results in increased ADP/ATP ratio. This low energetic state would be detrimental for viral replication. Since cells with time do restore the rest membrane potential, large conductance K^+^ channels (BK channels), that are both calcium and voltage-gated, would cause an outward K^+^ current, reestablishing rest membrane potential. Thus, the imbalance of the intracellular K^+^ concentration may also affect the functioning of different potassium channels that may be important for the virus life cycle. The illustration was created with the web-based tool BioRender (https://biorender.com).

The data we obtained about the membrane potential points to a possible explanation about the inhibition mechanism of SARS-CoV-2 infection in Vero cells by extracellular increased concentration of NaCl. Various pulmonary virus infections impair the activity of ENaC, and the increased activation state of these channels can significantly decrease the influenza A infection ^25–27^. Additionally, the spike and envelope small membrane protein (E) of SARS-CoV can directly decrease the inward current of ENaC as well as decrease its expression through the PKC signaling pathway^28^. Impairment of ENac activity itself would be enough to explain the pulmonary edema in the alveoli, since its inhibition would increase the amount of Na^+^ in the airway, and water would flow by osmosis ^29^.

Membrane potential has been shown to be important for some virus entry, and in the case of human Rhinovirus Type 2 infection the membrane hyperpolarization enhances infection^30^. Inhibition of membrane depolarization is part of viral lung infection strategy, as shown for SARS-CoV^21^ and respiratory syncytial virus ^31^, whose cell infection inhibits the entry of Na^+^ into the cell. While the ENac is voltage-insensitive, the gradient between high and low Na^+^ (out- and inside of the cell) forces the entry of Na^+^, being an unfavorable condition for SARS-CoV-2 infection.

Our results show that the increase of NaCl concentration causes an immediate membrane depolarization, probably through the activation of an extracellular Na^+^ sensitive channel, called Na_X_^32^. The activation of Na_x_ leads to Na^+^ inward flow, causing the depolarization, which in turn can open the voltage gated Ca^2+^ channels Cav2.1 that are expressed at the apical membrane of epithelial cells ^33^. Moreover, Na_x_ activation upregulates prostasin (protease) release into the extracellular space activating then ENaC by cleaving the extracellular loop of the γ ENaC subunit ^34–36^. Following Na^+^ influx via ENaC, the cells also present downstream mRNA synthesis elevation of inflammatory mediators ^36^. In the same line, Na_x_ expression blockade can improve scarring ^36^. Therefore, extracellular Na^+^ imbalance through ENac inefficiency by SARS-CoV-2 infection may explain the formation of pulmonary fibrosis due to virus infection

We observed mainly two patterns of depolarization: (1) lower doses of NaCl concentration triggered depolarization and after 180s the cell membrane reestablished the resting state, causing a timid inhibition of the virus release, while (2) doses higher than 210 mM could trigger a high amplitude depolarization, in which the cells needed periods longer than 24h to recover the resting state, causing over 90% inhibition of the virus. In the first situation, we hypothesize that activation of Ca^2+^ sensitive K^+^ channels and Na^+^/K^+^ ATPase pump could recover the steady state, while in the second situation, we hypothesize that the over stimulation of Na_x_, ENaC and Ca2.1, would overwhelm the cell, consume the ATP production, thus stressing the mitochondria. Two types of Ca^2+^-dependent K^+^ channels, small (SK) and big (BK) conductance, can be responsible for Ca^2+^-activated outward K^+^ currents. While SK channels are expressed especially in the central nervous system and heart, being responsible for the afterhyperpolarization of excitable cells, BK channels are broadly expressed in mammalian cells. These two K^+^ channel subtypes differ mainly in their single channel conductance, which is in the order of 10 pS for SK channels and 150-250 pS for BK channels ^37^. This large K^+^ conductance makes BK channels important for cell membrane repolarization, since BK channels are activated by membrane depolarization and/or high cytosolic Ca^2+^ concentration ([Ca^2+^]_i_). Literature reports that BK channels are activated in Vero cells during hypotonic stress, where cells are submitted to a 45% less NaCl environment, in which BK channels were activated by Ca^2+^ released by intracellular stores^38^. Here, membrane potential assays showed that Vero cells were able to cope with high NaCl concentrations and recover its resting membrane potential, probably due to activation of BK channels, which would activate either by the observed depolarization, or by the Ca^2+^ inward flow that probably occurred.

Our results showed that SARS-CoV-2 inhibition effects occurs at the onset of hypertonic stress, since the viral inhibition did not change when Vero cells had been only pre-treated with NaCl (**Fig 1a**, AD curve). Mitochondrial membrane potential can also be determinant for antiviral cell responses. In human embryonic kidney (HEK) and mouse embryonic kidney (MEF) cells, the application of mitochondrial uncoupler CCCP, which dissipates the transmembrane potential through the increase of membrane permeability, caused an impaired antiviral response by the cell ^39^. This did not occur for the F_1_F_O_ ATP synthase inhibitor oligomycin B, indicating that the diminished antiviral effect observed was associated with mitochondrial membrane depolarization, but not with impaired ATP synthesis alone. Our hypothesis shown in **Fig. 3 b** is that the overflow of Na^+^ in Vero cells, followed by increased [Ca^2+^]_i_, would cause an extreme low energy state (high ADP/ATP ratio), leading to impaired virus replication. Another possible effect is that the increased concentration of intracellular Na^+^ that causes membrane depolarization results in a decreased intracellular K^+^ concentration due to restoring the membrane potential. Thus, the imbalance of the intracellular K^+^ concentration can affect the functioning of different potassium channels that are important for the life cycle of some viruses, as in the case of HIV^40^.

It is known that hyperosmotic stress induced by only 30 mM above isotonic concentration, resulting in 150 mM NaCl, dissipates mitochondrial membrane potential within minutes, whereas the addition of 100 mM NaCl (220 mM NaCl final concentration) causes instant mitochondrial depolarization in Vero cells, with no detectable cytochrome c release to the cytoplasm^41^. In addition, high intracellular Na^+^ concentrations stimulate ion active transport through Na^+^/K^+^ ATPase, and an hyperosmotic stress caused by NaCl (and not sorbitol or mannitol) causes an increase of the pump activity, since elevated chloride levels specifically increases the expression of the Na^+^/K^+^ ATPase γ-subunit ^42^.

In our study, mitochondrial depolarization combined with the elevated activity of this ATP-consuming pump probably resulted in an extremely low energetic state, with increased ADP/ATP ratio, which would compromise viral replication. Besides the energetic state, mitochondria are associated with other cellular functions that can be interesting for viral infection. In A549 cells, a cell line often used as a model of type II pulmonary epithelial cells, the SARS-CoV ORF-9b protein localizes to mitochondria and reduces the levels of DRP1, a GTPase that regulates mitochondrial fission. This resulted in enhanced fusion and elongated mitochondria compared to control cells, which led to an impaired innate immune system signaling, necessary for viral replication^43^. Last, higher [Ca^2+^]_i_ levels can cause damage not only in mitochondria, but also for viral proteins. High levels of [Ca^2+^]_i_ can, besides triggering membrane depolarization on Vero cells, disturb [Ca^2+^]_i_ levels, which can lead to impaired viral replication. There is solid evidence that viral infections and the expression of viral proteins modify [Ca^2+^]_i_, since Ca^2+^ is an important second messenger and regulator of many cellular processes^44^. In virus-host interactions, viruses can use different calcium-binding proteins of different cell compartments, including cytoplasm and endoplasmic reticulum, such as calmodulin, proteins of the S100 family and calnexin, and a disturbance in these metabolic processes would be detrimental for viral replication.

Since NaCl decreases viral replication by more than 90%, probably due to a mechanism associated with the effect of membrane depolarization, our results suggest that tests on humans should be carried out to validate the effectiveness of hypertonic solution treatment in patients with COVID-19. This treatment could be effective for either those, who are hospitalized, or as prophylaxis treatment for non-hospitalized cases. It is worth to mention that this procedure is already used to improve lung function in cystic fibrosis patients^45^ and nebulized 3% hypertonic saline treatment for infants with moderate to severe bronchiolitis is safe without adverse events such as bronchospasm, cough or wheezing aggravation^46–48^. However, this safe protocol has not, to our knowledge, been used to treat COVID-19 patients. The establishment of a therapeutic role for hypertonic saline solution in SARS-CoV-2 infection has relevant clinical implications: it may significantly reduce the duration of hospital internment and improve the clinical severity score. This inexpensive therapy should be tested in COVID-19 patients. We have developed a website, www.icbcorona.com (see Supplementary Material for more information), in which physicians can input and share their observations of clinical cases, in which patients were treated with hypertonic solution (inhalation with 1.5% NaCl or greater). In this way, protocols can be improved, and the evolution of their patients can be shared with the scientific community. Furthermore, findings from a post-hoc analysis of NCT02438579 clinical trial suggest that hypertonic saline nasal irrigation and gargling may have play a role in reducing symptoms and duration of illness caused by COVID-19^49^. In addition, there are three clinical trials registered in ongoing status (NCT04465604, NCT04382131 and NCT04341688) according to ClinicalTrials.gov (https://clinicaltrials.gov/ct2/home), related to the use of hypertonic saline solution for COVID-19 treatment. This scenario highlights the importance of our results and reinforces the need of carrying out the current clinical studies for a better understanding of the benefits of hypertonic saline treatment for COVID-19 patients.

In summary, hypertonic saline solution inhibits SARS-CoV-2 virus replication in Vero cells due to perturbation in one or several steps of the virus intracellular cycle. NaCl is known to drive type I interferon signalling in macrophages, and that elevated concentrations of NaCl enhance the production of HOCl in non-myeloid cells^14,50^. Hypertonic saline solution decreased Respiratory syncytial virus infection and pro-inflammatory response such as IL-6 and IL-8 release in cultures of human respiratory epithelial lines^51^. Therefore, the complete molecular mechanism underlying the observed phenomenon may be more complex, involving several factors that together result in the inhibition of viral infection. Therefore, further assays must be done to elucidate the real mechanism of virus inhibition by high concentrations of NaCl.

## Methods

### Cell, virus and NaCl dilution

African green monkey kidney Vero cell line (ATCC CCL-81) were maintained in Dulbecco’s modified eagle medium (DMEM), supplemented with 10% fetal bovine serum (FBS), 1% nonessential amino acids (NEAA), 1% sodium pyruvate (Sigma-Aldrich Co., Deisenhofen, Germany) and incubated in a humidified atmosphere containing 5% CO_2_ at 37°C. A clinical isolate SARS-CoV-2/SP02/human/2020/BRA (GenBank access n° MT126808.1) was propagated in Vero E6 cells, and viral titer was determined by 50% tissue culture infective dose (TCID_50_) and by plaque forming units per milliliter (PFU/mL), as previously described by Araujo and collaborators^52^. We conducted all the infection experiments in a biosafety level-3 (BLS-3) laboratory at the Institute of Biomedical Sciences, University of São Paulo, São Paulo, Brazil, following the Laboratory biosafety guidance related to the novel coronavirus (2019-nCoV) by WHO^53^. Sodium chloride (NaCl) was diluted in free-DMEM that already contains in its composition 110 mM of NaCl. In that way we assumed seven plus additional increasing concentrations (25, 50, 75, 100, 125, 150 and 175 mM) of NaCl in the cell medium resulting in a final NaCl concentration of 135, 160, 185, 210, 235, 260 and 285 mM.

### Cell viability assays

For cell viability and stress, 5×10^4^ cells on black clear bottom 96 well plates stained with the AlamarBlue™ Cell Viability Reagent (Thermo Fisher Scientific), according to manufacturer’s instructions. Cells were stained and fluorescence intensity recorded at rest (point 0), 1h, 24hs and 72hs after NaCl challenge, using 530 nm for excitation and 560/590 for emission detection in the FlexStation III microplate reader. Data were normalized by a blank control that consists of cell medium for natural reduction conditions. Data are plotted as reference to the basal condition of 110 mM NaCl concentration (DMEM’s salt concentration). Cell viability was further evaluated using a colorimetric assay by quantifying lactate dehydrogenase (LDH) released into the culture supernatant from cells with damaged membranes, using the CytoTox 96® Non-Radioactive Cytotoxicity Assay (Promega Corp., Madison, WI). Detection was performed using a microplate reader (POLARstar^®^ Omega, BMG LABTECH, Ortenberg, Germany) at 492 nm. The activity of the released LDH was reported as a percentage of the total cellular LDH (measured after the complete lysis of control cells corresponding at the maximal amount that can be released by cells, therefore 100%). Cell viability was normalised to untreated cells (0 mM NaCl added). Viability below 70% was used as a threshold for cytotoxicity ^54^.

### Cell death quantification

For cell death detection, we adapted a usual method for flow cytometry, based on staining live cells with the membrane impermeant dye, propidium iodide (PI), and the permeable fluorophore, Hoechst 33342. Since PI stains the dead cells, while Hoechst 33342 stains every cell in the dish, cell death rates were quantified by the total fluorescence intensity of the PI staining in the well, normalized by the total fluorescence intensity of the Hoechst 33342 ^55^. The recordings were acquired at rest (point 0h), 1h, 24hs and 72hs after NaCl challenge, using 535 nm for excitation and 617 nm for emission detection for PI, and 350 nm for excitation and 461 nm for emission of Hoechst 33342 in the FlexStation III microplate reader. For positive control of total cell death, cells were lysed with 1% TritonX-100 detergent.

### Nucleic acid extraction and quantitative real-time RT-PCR (RT-qPCR)

In order to perform the quantification of SARS-CoV-2 viral load, the extraction of total nucleic acid (RNA and DNA) from the collected cell culture supernatant was carried out, using the semi-automated NucliSENS® easyMag® platform (BioMerieux, Lyon, France), following the manufacturer’s’ instructions. The quantification of viral RNA was done using the AgPath-ID One-Step RT-PCR Kit (Applied Biosystems, Weiterstadt, Germany) on an ABI 7500 SDS real-time PCR machine (Applied Biosystems) using a reference published sequence of primers and probe for E gene ^56^. Numbers of RNA copies/mL were quantified using a specific in vitro-transcribed RNA quantification standard, kindly granted by Christian Drosten, Charité - Universita□tsmedizin Berlin, Germany, as described previously ^57^.

### Antiviral activity of NaCl during different time-of-addition

Vero CCL-81 cells were seeded in a clear-bottomed 96-well plate (5 × 10^4^ cells/mL) and incubated for 24h at 37°C for cell adherence. Then, they were treated with increasing concentrations of NaCl at different stages of virus infection as described below. Four different NaCl time-of-addition were evaluated^14,58^, compromising virus pre-incubation (VPI), absorption (AD), post-infection (PI) and adsorption plus post-infection, named full-time (FT). For the VPI treatment SARS-CoV-2 was pre-incubated with the increasing concentrations of NaCl for 1 h before infecting the cells. After adsorption, the inoculum was removed, replaced with media and maintained till the end of the experiment. For AD treatment, the different concentrations of NaCl were added to the cells monolayer for 1 h prior to virus infection, and maintained during 1 h for viral attachment process. Then, the virus-NaCl mixture was replaced with fresh DMEM until the end of the experiments. During the PI-staining experiment, the virus was added to the cells to allow infection for 1 hour, and then virus-containing supernatant was replaced with different NaCl concentration-containing medium until the end of the experiment. For FT treatment, Vero CCL-81 cells were pretreated with different concentrations of NaCl for 1 h prior to virus infection, followed by incubation with virus for 1 h in the presence of the NaCl. Then, the virus mixture was removed, cells were and cultured with the same concentrations of NaCl-containing medium until the end of the experiment. For all experimental groups, cells were infected with virus at a multiplicity of infection (MOI) of 0.02, and at 72 h.p.i.. Then cell supernatants were collected for RT-qPCR, and cells were screened for the presence/absence of cytopathic effects (CPE) under an optical microscope (Olympus, Tokyo, Japan).

### Analysis of membrane potential variation by microfluorimetry

Changes of the membrane potential of Vero cells exposed to increasing concentrations of NaCl was determined by plate microfluorimetry recordings with the FlexStation III microplate reader and the FLIPR Membrane Potential Assay Kit (Molecular Devices Corp., Sunnyvale, CA) following the manufactures’ instructions, as previously described^59^. The kit provides results in good correlation with those obtained in patch-clamping assays. Recordings of the fluorescence intensity of 5×10^4^ cells in black clear bottom of 96 wells plates were acquired at rest (point 0h), 1h, 24hs and 72hs after NaCl challenge (depolarizing agent). Time kinetics were obtained by measuring at 1.52s intervals for 120 s after 30 s of monitoring basal fluorescence intensity. Responses were calculated as the peak fluorescence minus the basal fluorescence. Fluorescence intensity was determined using the SoftMax2Pro software (Molecular Devices Corp.).

### Statistical analysis

Data were expressed as mean values ± SEM (standard error of the mean) of three independent experiments, with each measure performed in triplicate. The percentage (%) of inhibition and cytotoxicity were calculated and normalized to untreated (110 mM NaCl) cells. *p* values were calculated using unpaired two-tailed t-test or one-way ANOVA, when appropriate. *p* ≤ 0.05 was considered significant. The graphics were designed using GraphPad Prism 8.4.1 software.

## Acknowledgements

This work was supported by grants of the São Paulo Research Foundation (FAPESP Projects No. 2018/07366-4, 2019/00195-2, 2020/04680-0, 2017/24769-2, 2020/06409-1, 2016/20045-7 and 2020/06409-1) and Coordenação de Aperfeiçoamento de Pessoal de Nível Superior (CAPES number 88887.131387/2016-00).

