## supplemental information for "Hypertonic saline solution inhibits SARS-CoV-2 in vitro assay"

**Website development.** The website was developed by MAXXIDATA, a partner company, and the questions were divided in three blocks, all of them available in a logged area: [1] Clinical Case; [2] Treatment; [3] Results (**Figure S1**). The goal is to have this form filled after the treatment is finished, and final results are already available. In the clinical case block, the questions are related to the initial state of the patient, with no personal identification. Below are the questions in the website, for each block:

1. Questions for <u>[1] clinical case</u> block: (1.a). Is positive tested to COVID-19?; (1.b) Days since 1$^{st}$ symptoms to treatment, (1.c). Patient was intubated (yes or no); (1.d). Severity of this case (Asymptomatic, Mild or Severe); (1.e) Gender of the patient; (1.f) Initials or your own internal patient ID (this is for their internal control, and it is asked they should not identify the patient); (1.g) Data of birth; (h). Did the patient have any comorbid?
2. Questions for <u>[2] treatment</u> block: (2.a). Type of application (Puff, Inhaler, Nebulizer or other); (2.b) Hypertonic concentration; (2.c) Volume in mL; (2.c) Application frequency (every how many hours); (2.d) What else did you use? (chloroquine, azithromycin, hydroxychloroquine and/or other).
3. Questions for <u>[3] Results</u> block: (3.a). Treatment outcome: (from 0% to 100%); (3.b) After how many days did you see this result? (3.c) Area for the user to add comments about the case.

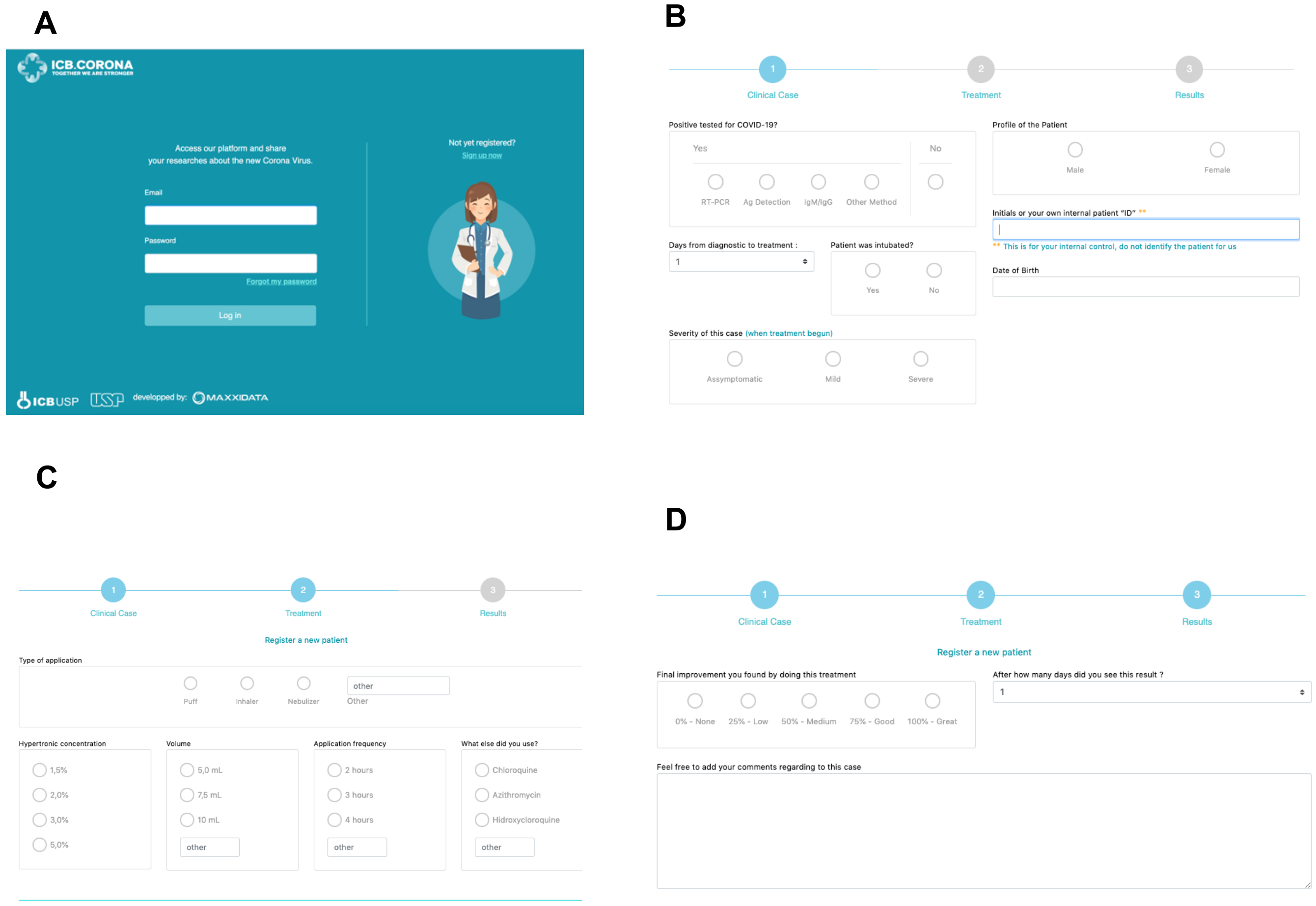


**Figure S1:** The website www.icbcorona.com. Physicians can input and share their observations of clinical cases in which patients were treated with hypertonic solution (inhalation with 1,5% NaCl or greater). The Login (**A**), Clinical Case (**B**), Treatment (**C**), and Results blocks (**D**) are shown.
